# Molecular basis for voltage sensitivity in membrane proteins

**DOI:** 10.1101/295816

**Authors:** Marina A. Kasimova, Erik Lindahl, Lucie Delemotte

## Abstract

Voltage-sensitive membrane proteins are united by the ability to transform changes in the membrane potential into mechanical work. They are responsible for a spectrum of key physiological processes in living organisms, including electric signaling and progression along the cell cycle. While the voltage-sensing mechanism has been well characterized for some membrane proteins such as voltage-gated ion channels, for others even the location of the voltage-sensing elements remains unknown. The detection of these elements using experimental techniques is complicated due to the large diversity of membrane proteins. Here, we suggest a computational approach to predict voltage-sensing elements in any membrane protein independent of structure or function. It relies on the estimation of the capacity of a protein to respond to changes in the membrane potential. We first show how this property correlates well with voltage sensitivity by applying our approach to a set of membrane proteins including voltage-sensitive and voltage-insensitive ones. We further show that it correctly identifies true voltage-sensitive residues in the voltage sensor domain of voltage-gated ion channels. Finally, we investigate six membrane proteins for which the voltage-sensing elements have not yet been characterized and identify residues and ions potentially involved in the response to voltage. The suggested approach is fast and simple and allows for characterization of voltage sensitivity that goes beyond mere identification of charges. We anticipate that its application prior to mutagenesis experiments will allow for significant reduction of the number of potential voltage-sensitive elements to be tested.

## Introduction

The membrane potential (MP) in living cells results from the uneven distribution of ions between the two sides of the cell membrane (1). It regulates several critical physiological processes such as the formation and propagation of the action potential in excitable cells (1) and the progression along the cell cycle in non-excitable ones (2–4). Voltage-sensitive membrane proteins are able to detect changes in the MP and in some cases to act directly on it (5). The ability to understand and eventually to modulate the function of these proteins provides an interesting strategy to diagnose and treat neurological diseases by changing either the MP or the proteins’ response to it.

The group of voltage-sensitive membrane proteins (VSMPs) includes representatives from transporters (5–37), enzymes (38–44), receptors (45, 46) and proteins playing a role in structure or adhesion (47) (Fig. 1). These proteins are extremely diverse in their structure, but all of them have a common ability to convert the electrical energy into a mechanical response. To achieve this, VSMPs have developed one or more voltage-sensing elements, which detect changes in the MP and alter their conformational state accordingly (5, 48). In voltage-gated potassium and sodium channels, the protein itself plays the role of the voltage sensor: the S4 helix of the S1-S4 helical bundle carries several positively charged residues, which upon application of an electric field move in the direction of this field (49–59). In most other VSMPs, the voltage-sensing elements are nowhere near as well characterized. Several studies suggest different origins of their voltage sensitivity, including specific protein residues and/or ions trapped inside protein cavities (8, 9, 60–67). Ions moving in an electric field have for instance been suggested to trigger conformational rearrangements in voltage-gated chloride channels (66, 68–70), solute carriers (10), active transporters (8, 9, 71, 72) and G-protein coupled receptors (67).

**Figure 1:**
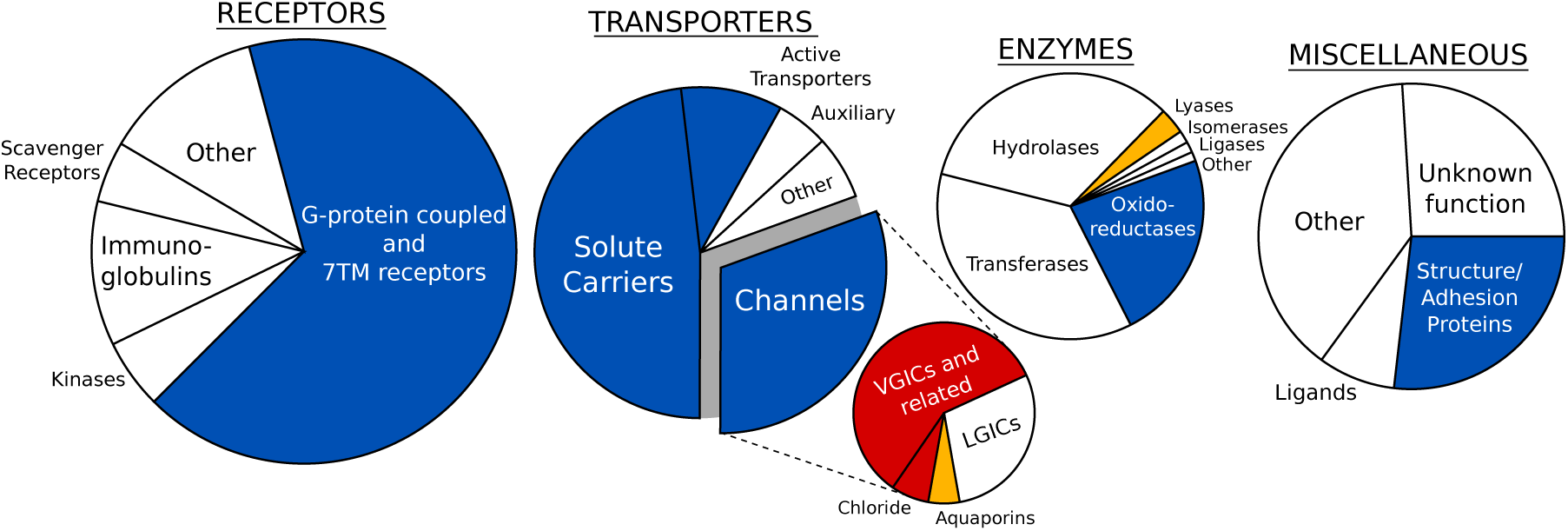
Membrane protein families, encompassing voltage-sensitive representatives. The families are arranged in four large groups according to (73) (only α-helical membrane proteins are shown). Channels are subdivided in voltage-gated (VGIC), ligand-gated (LGICs), Chloride channels and Aquaporins. Blue indicates families containing voltage-sensitive membrane proteins, and white indicates families in which voltage-sensitive representatives were not discovered so far. The families colored in red correspond to membrane proteins for which changes in the MP is the primary stimulus for activation. Finally, we colored in orange the families containing membrane proteins whose voltage sensitivity is controversial. To compose this figure we used data from references (5–47).

In this work, we characterize the voltage sensitivity of several selected membrane proteins using a newly developed computational approach. It estimates how fast a local electric field acting on a protein changes upon application of an external electric field; we will call this property the local electric field response. It also estimates the system’s response capacity, which reflects the ability of the system to detect changes in a local electric field and to respond to them. The system’s response capacity has a direct connection to the gating charge, a well-known characteristic of voltage sensitivity that corresponds to the amount of charge transferred during protein activation. We show that in all tested voltage-sensitive proteins the system’s response capacity is large compared to voltage-insensitive ones, which suggests this could be an efficient tool to probe voltage sensitivity in other membrane proteins. We also show how the local electric field response can be large in selected regions of voltage-insensitive proteins. However, the lack of charges in these regions results in their inability to respond to changes in the MP. Finally, we investigate six VSMPs for which the voltage sensors have not yet been characterized. For each of them, we depict putative voltage-sensitive residues and/or voltage-sensitive ions trapped in protein cavities.

Our approach only requires knowledge of the protein structure and modest computational resources, which makes it possible to apply prior to mutagenesis experiments in order to reduce the number of potential voltage-sensitive elements to be tested.

## Results and Discussion

### In voltage-sensitive membrane proteins charges are located in regions with a large local electric field response

Voltage sensitivity occurs in many different membrane proteins with diverse structure and function (5–47) – the unifying factor is how they are all able to detect changes in the membrane potential (MP) and to convert these changes into mechanical work (5). Our approach to characterize voltage-sensing elements is based on the hypothesis that the force acting on these elements increases substantially upon an increase in the MP. Thus, they are located in regions where a change in the MP *∂V*_*m*_ results in the largest change in the local electric field *∂E* (*r*). We define the rate of change of *E* (*r*) with respect to *V*_*m*_ as the local electric field response *R* (*r*):

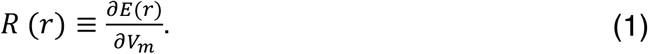

The actual system’s response, on the other hand, depends on the interplay between *R* (*r*) and the system charges *q*_*i*_ = {*q*_1_, …, *q*_*N*_}. Here, *q*_*i*_ plays the role of an effector translating changes in the local electric field into mechanical work. We define the capacity of the system residue *j* to respond to changes in the local electric field *C*^*j*^ as:

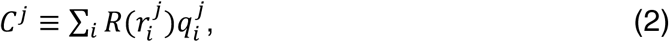

where the summation runs over all charges *q*_*i*_ of the residue *j. C*^*j*^ has a direct connection to the gating charge *Q*^*j*^ it corresponds to the gradient of *Q*^*j*^ along the direction of the external electric field *z* (cf. Methods):

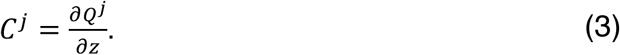

*Q*^*j*^ is a characteristic variable of voltage sensitivity, which was first proposed to report the amount of charge transferred by a residue *j* upon gating in voltage-gated ion channels (74–76). It can also be generalized to any activation process rather than only gating and therefore be applied to any given voltage-sensitive membrane protein. *C*^*j*^, on the other hand, reports the amount of charge transferred by a residue *j* upon 1 Å displacement (along *z*). Therefore, *Q*^*j*^ can be viewed as a summation of *C*^*j*^ for all protein conformations involved in the activation process. Since in many cases this process remains largely uncharacterized, *C*^*j*^ of a single protein conformation can be considered as a good predictor for *Q*^*j*^.

Our approach calculates, first, the local electric field response *R*, depicting the system’s regions subject to voltage sensitivity, and second, the system’s response capacity *C*^*j*^, depicting the residues with the potential to contribute to the gating charge.

To test the *R*&*C*^*j*^ approach, we applied it to a set of voltage-sensitive and voltage-insensitive membrane proteins. In the former case, we have considered two voltage-gated ion channels, whose voltage-sensing elements have been well characterized, and six other membrane proteins, for which these elements remain largely unknown (see Table 1).

**Table 1.**
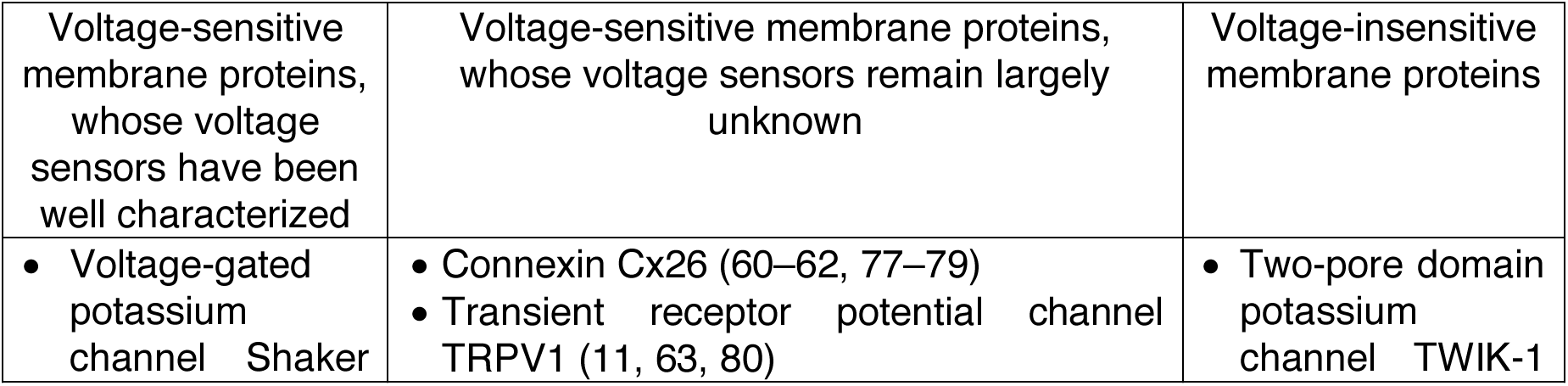
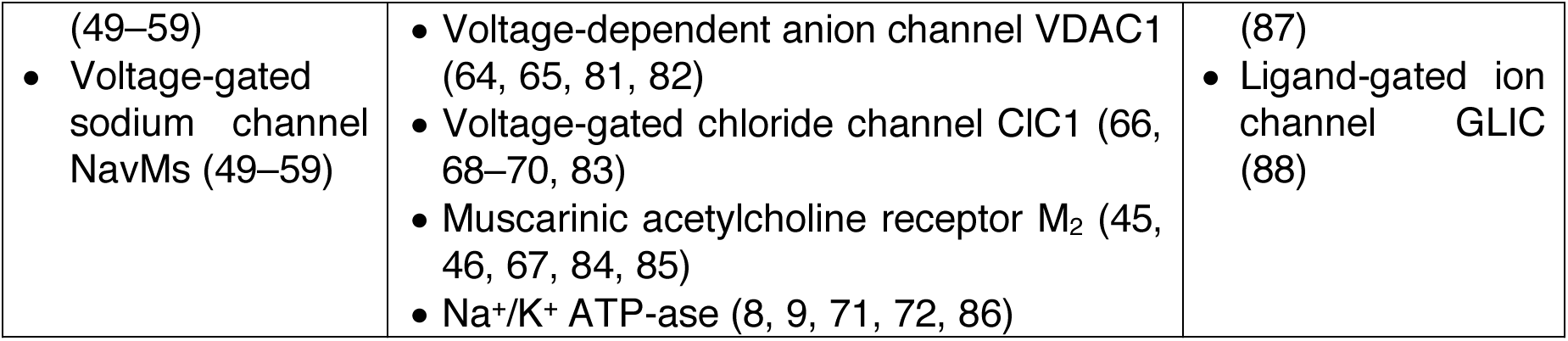

For each protein, we calculated the local electric field response *R* and the system’s response capacity *C*^*j*^ (Fig. 2). In many voltage-sensitive membrane proteins, we detected regions with large *R* and therefore with a potential to respond to changes in *V*_*m*_. The center of the voltage sensors in Shaker and NavMs (49–59), the N-terminus in VDAC1 (64, 65, 81, 82), and the Cl-binding site in ClC1 (66, 68–70, 83) all show *R* larger than 0.08 Å^-1^. However, in the M_2_ receptor (45, 46, 67, 84, 85), which is also known to be voltage-sensitive, *R* is small and reaches a maximum of only 0.04 Å^-1^ in the Na^+^ binding site. Moreover, in voltage-insensitive membrane proteins, large *R* values were detected along the conductive pore, with the selectivity filter in TWIK-1 (87) and the gate in GLIC (88) showing *R* as large as 0.08 Å^-1^. Therefore, *R* does not directly correlate with voltage sensitivity and cannot be used to discriminate between voltage-sensitive and voltage-insensitive proteins.

**Figure 2:**
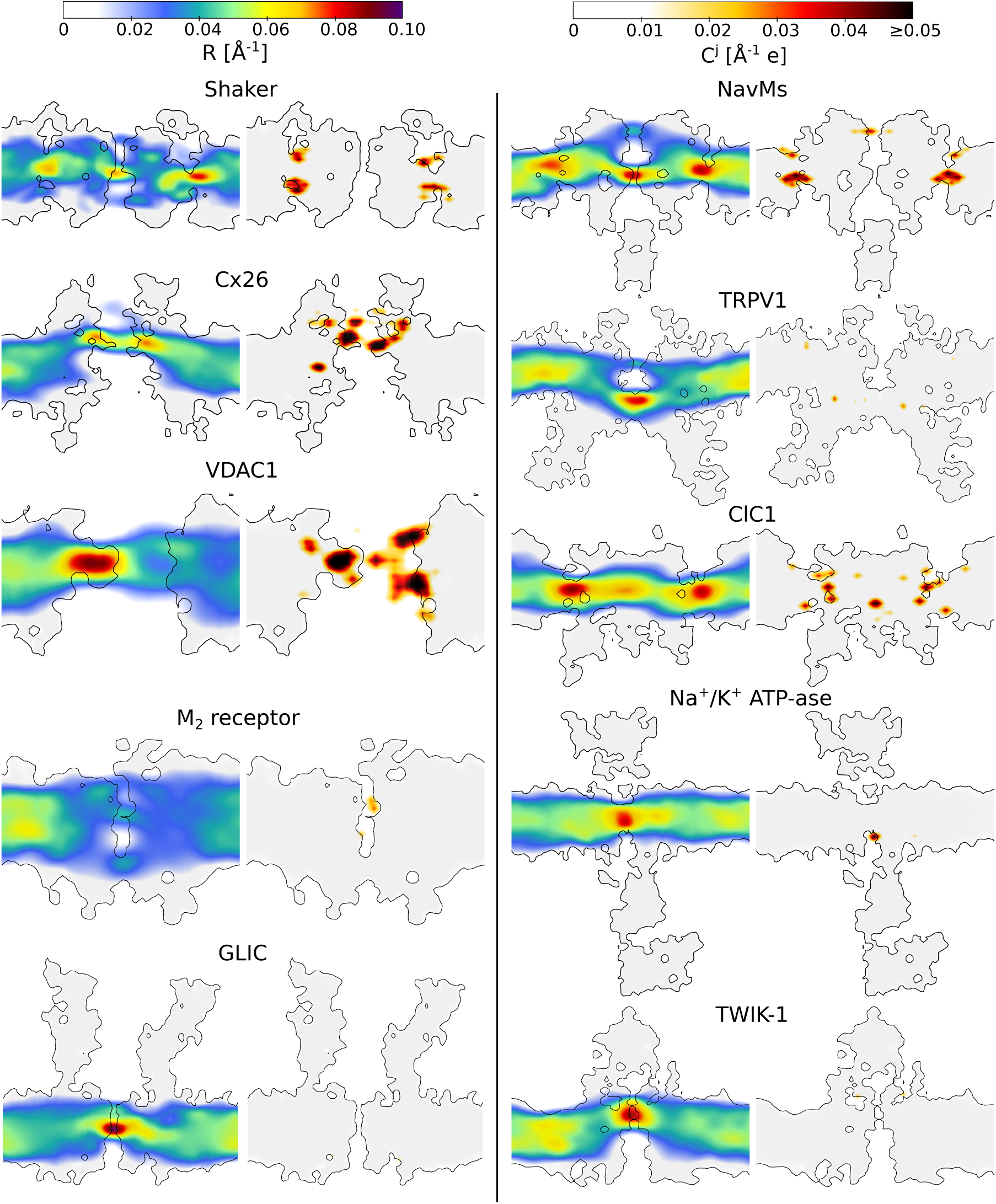
The local electric field response *R* (left) and the system’s response capacity *C*^*j*^ (right) estimated for eight different membrane proteins: voltage-gated potassium channel Shaker (89, 90), voltage-gated sodium channel NavMs (91), connexin Cx26 (92), transient receptor potential channel TRPV1 (93–95), voltage-dependent anion channel VDAC1 (96), voltage-gated chloride channel ClC1 (97), muscarinic acetylcholine receptor M_2_ (98), Na^+^/K^+^ ATP-ase (99), ligand-gated ion channel GLIC (100) and two-pore domain potassium channel TWIK-1 (101). The slices of the systems along the normal to the membrane are shown. The grey area shows the regions, which are not accessible to water, *i.e.* the proteins and the membrane. In order to clearly represent *C*^*j*^, we approximated each point charge of the system element *j* with a Gaussian distribution (σ = 1.5 Å) and then integrated the signal over 25 slices (each 1 Å width) parallel to the plane shown in the figure. Only the values above the detection threshold were considered for the integration (cf. Methods). Shaker, NavMs, Cx26, VDAC1 and ClC1, for which changes in the MP is the primary stimulus for activation, have the largest *C*^*j*^, while TRPV1, which is known to be very weakly voltage-sensitive (11, 63, 80), has the smallest *C*^*j*^ among the voltage-sensitive membrane proteins.

*C*^*j*^, on the other hand, correlates rather well with voltage sensitivity: for all voltage-sensitive membrane proteins *C*^*j*^ is large, while it is close to or below the detection threshold for all voltage-insensitive ones. Together with the previous observation, this suggests that voltage-sensitive proteins have evolved to place their charges in the regions with large *R* and therefore maximize their potential to respond to changes in *V*_*m*_. In contrast, in voltage-insensitive proteins, charges are located far from the regions with large *R* and thus do not sense changes in the local electric field. This also suggests our approach can be used to probe voltage sensitivity in other membrane proteins, in which this property has not yet been tested.

### In voltage-gated ion channels the residues with large *C*^*j*^ correspond to the true voltage sensors

We further describe the results obtained for the two voltage-gated ion channels, Shaker and NavMs. For voltage-gated ion channels in general, the mechanism of voltage sensitivity has been well characterized based on numerous experimental and computational studies (49–59). These channels have four-helix bundle domains that sense changes in the MP – voltage sensor domains. One of the four helices (S4) carries several positively charged residues, while the other three (S1-S3) include a few negative countercharges (50, 51). Upon application of an electric field, the positive charges of S4 are displaced along the direction of this field to trigger conformational rearrangements in the pore domain (49, 52–58).

Our approach correctly detects the positively charged residues on S4 and their negative countercharges on S1-S3 as voltage-sensing elements (Fig. 3). Among them, residues located in the center of the voltage sensor domain show large *C*^*j*^, while those exposed to the extracellular or cytosolic solutions show *C*^*j*^ close to the detection threshold (cf. Methods). This indicates that, in the conformational states of Shaker and NavMs chosen for the analysis, residues such as R4 and K5/R5 contribute most to the gating charge, while the contribution from R1 and R2 is small.

**Figure 3:**
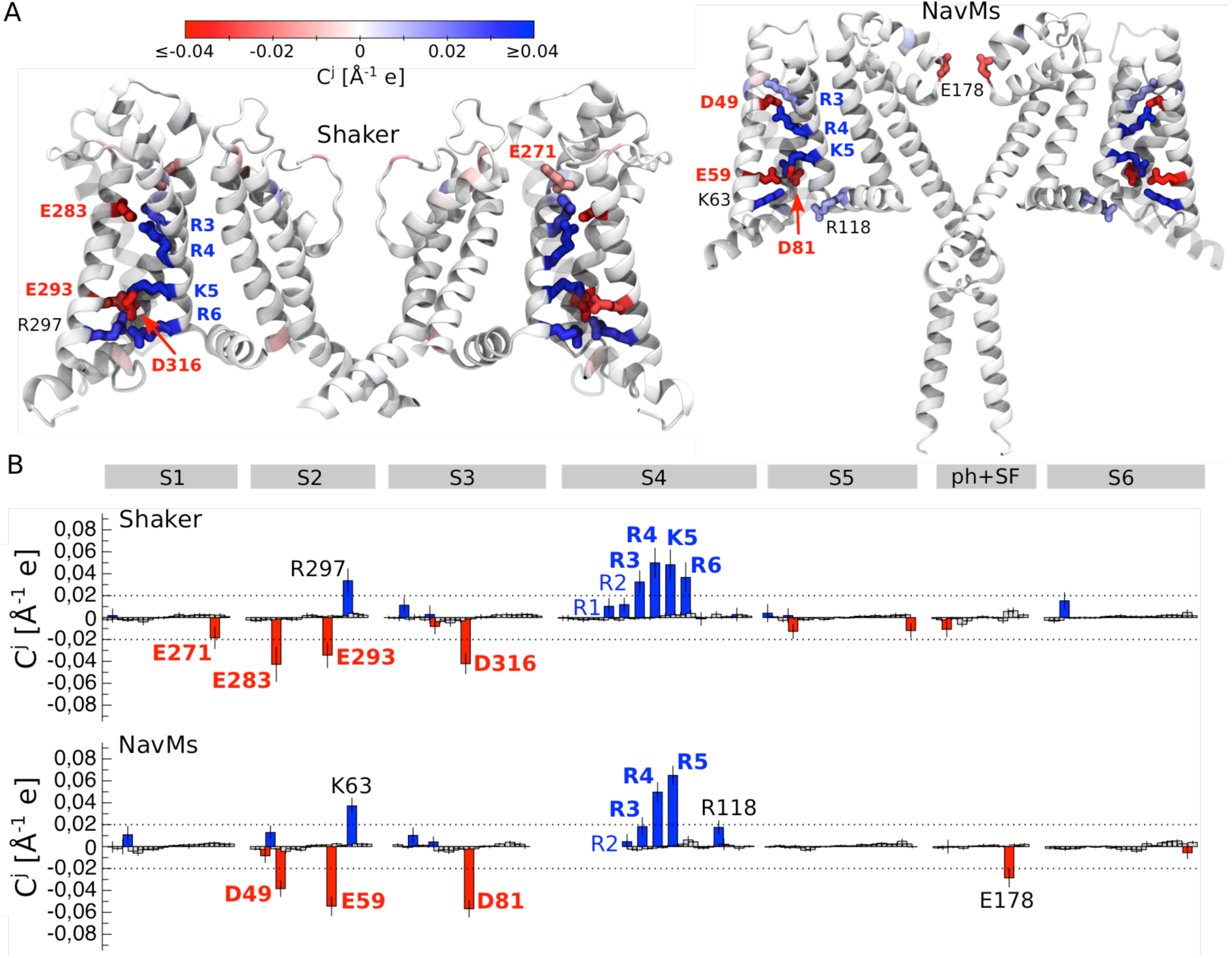
Detection of the voltage-sensing elements in the two voltage-gated ion channels, Shaker (89, 90) and NavMs (91). A. Cartoon representation of the two channels. The residues whose *C*^*j*^ is larger than the detection threshold are shown. B. *C*^*j*^ estimated for Shaker and NavMs; the average and the standard deviation are shown. The positively and negatively charged residues are shown as blue and red bars, respectively. The dashed lines represent the detection threshold. S1-S6 denotes the transmembrane segments, and ph+SF – the pore helix and the selectivity filter. The residues colored in blue or red and shown in bold correspond to the known voltage sensors and were detected by our method; those colored in blue and are not shown in bold – to the known voltage sensors, for which our method showed *C*^*j*^ below the detection threshold. Finally, residues colored in black were detected by our method but were not yet shown to play a role in voltage sensitivity.

A few other residues show *C*^*j*^ above the detection threshold, including R118 on the S4-S5 linker and E178 in the selectivity filter of NavMs, and a positive residue on S2 in both Shaker and NavMs. To our knowledge, the role of these residues in voltage sensitivity has not yet been assessed, and thus experimental verification will be required to confirm whether they are true or false positive signals.

### Estimation of the system’s response capacity allows for detection of voltage sensors in uncharacterized voltage-sensitive membrane proteins

The correct identification of voltage sensors in Shaker and NavMs strengthened our confidence in the predictive ability of our approach and suggested applying it to other membrane proteins, whose voltage sensors remain unknown. Overall, we applied it to six membrane proteins, including Connexin Cx26 (92), Transient Receptor Potential channel TRPV1 (93–95), voltage-dependent anion channel VDAC1 (96), voltage-gated chloride channel ClC1 (97), muscarinic acetylcholine receptor M_2_ (98), and Na^+^/K^+^ ATP-ase (99) (Fig. 4 and 5).

**Figure 4:**
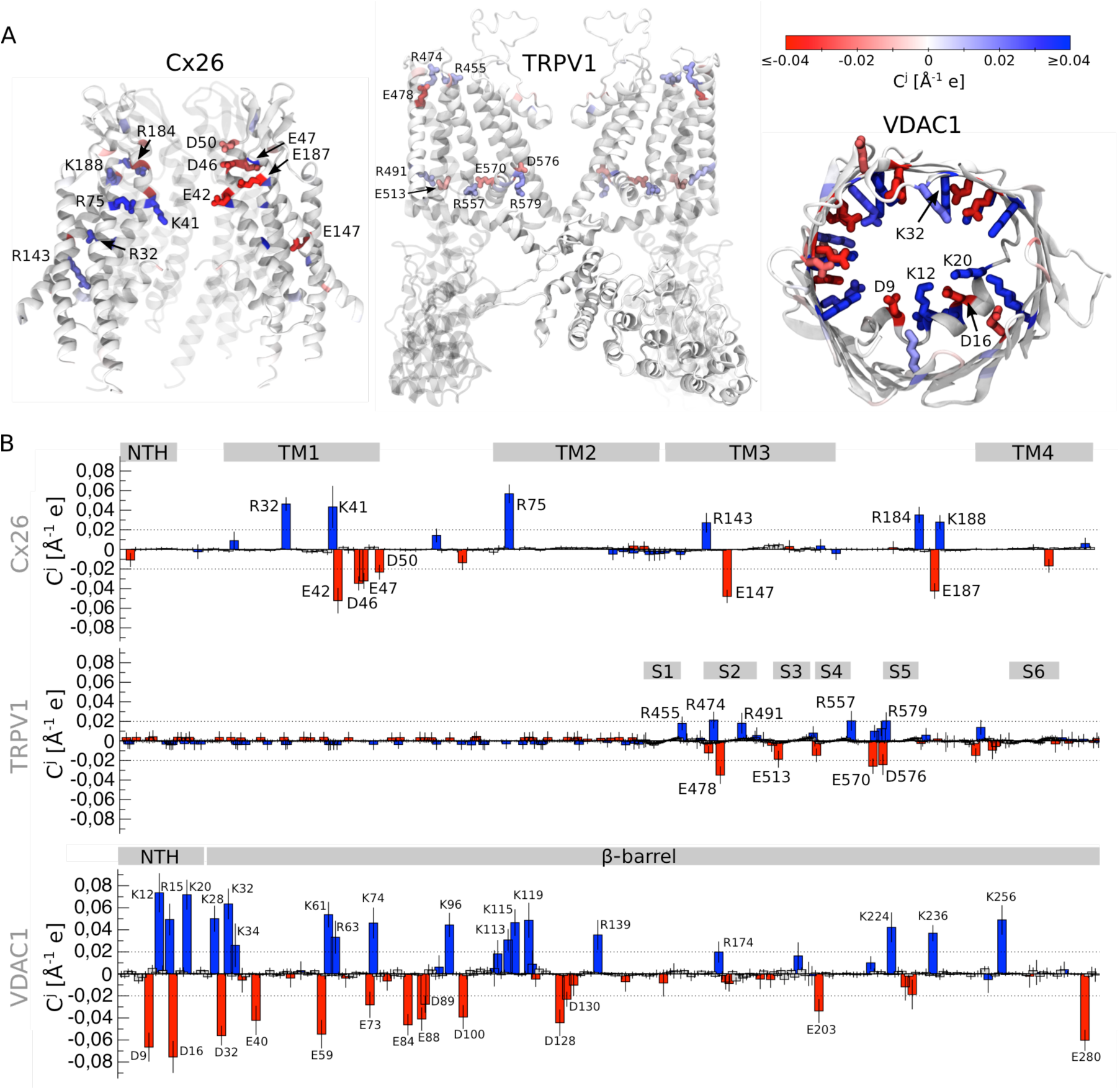
Detection of the voltage-sensing sensing elements in connexin Cx26 (92), transient receptor potential channel TRPV1 (93–95) and voltage-dependent anion channels VDAC1 (96). A. Cartoon representation of the membrane proteins. The residues whose *C*^*j*^ is larger than the detection threshold are shown. B. *C*^*j*^ estimated for Cx26, TRPV1 and VDAC1; the average and standard deviation are shown. The grey rectangles correspond to different regions of the proteins: NTH denotes the α-helical N-terminus in Cx26 and VDAC1, TM1-TM4 – transmembrane segments in Cx26, S1-S6 – transmembrane segments in TRPV1, and β-barrel – the β-barrel in VDAC1.

**Figure 5.**
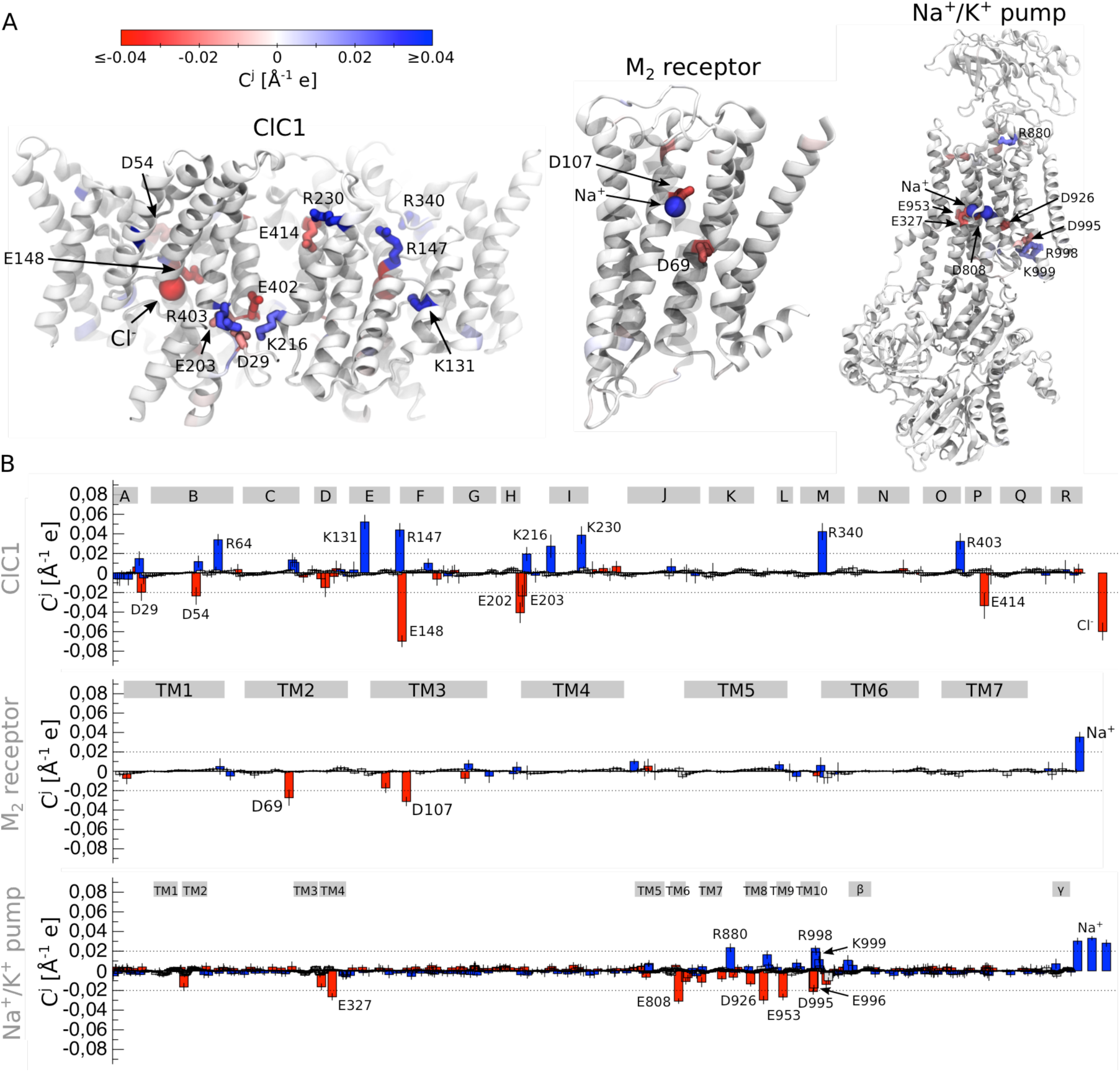
Detection of the voltage-sensing elements in the voltage-gated chloride channel ClC1 (97), muscarinic acetylcholine receptor M_2_ (98) and Na^+^/K^+^ ATP-ase (99). A. Cartoon representation of the membrane proteins. The residues whose *C*^*j*^ is larger than the detection threshold are shown. B. *C*^*j*^ estimated for ClC1, M_2_ receptor and Na^+^/K^+^ ATP-ase; the average and standard deviation are shown. The grey rectangles correspond to different regions of the proteins: A-R denotes the α-helical transmembrane segments in ClC1, TM1-TM7 – the transmembrane segments in M_2_ receptor, TM1-TM10 – the transmembrane segments in Na^+^/K^+^ ATP-ase, β and γ – auxiliary subunits of Na^+^/K^+^ ATP-ase.

For Cx26, we detected two regions with large *C*^*j*^, including the extracellular opening (E42, D46, D50, K41, R75, R184 and K188), and the center of the protein subunit (E147, R32 and R143) (Fig. 4). Many of these residues have already been suggested to play crucial roles in voltage sensitivity (60, 62, 77–79). For instance, molecular dynamics simulations (62, 79) suggested an electrostatic network between E42, D46, R75, R184, E187 and K188 to be a part of the voltage sensor. Experimental evidence indicates mutagenesis of R75, K41 and E42 significantly alters Cx26 voltage sensitivity (60, 77, 78). In particular, K41 neutralization results in an increase of the gating charge, while the double mutant K41E/E42S was shown to be more sensitive to voltage than the wild type (60). While experimental evidence suggests D2 to also contribute to voltage sensitivity (61), this residue did not exhibit any large *C*^*j*^ value in our analysis. We believe this discrepancy might be caused by the low resolution of the N-terminus, where D2 is located (92): shortly after the start of molecular dynamics simulations the N-terminus lost its secondary structure and significantly deviated from its initial position, which limits the predictive power in this region.

TRPV1 has a four-helical domain, which is similar to the voltage sensor domain of Shaker and NavMs (93, 94). However, since this channel does not include several crucial charges on S4 and the corresponding countercharges on S1-S3 (93, 94), it likely responds to changes in the MP through a different mechanism. Our analysis reveals that in addition to the lack of charges, the four-helical domain also shows small *R*, emphasizing that this region is not able to sense changes in the MP (Fig. 2). On the other hand, we found *C*^*j*^ values above the detection threshold for residues of the extra- and intracellular openings of the four-helical domain (R455, R474, E478, R491, E513) and of the S4-S5 linker (R557, E570, D576, R579) (Fig. 4). Mutagenesis of several of these residues (R557, E570, D576 and R579) has already been shown to significantly affect TRPV1 voltage sensitivity (63), which agrees well with our predictions.

In VDAC1, we observed multiple residues spread over the entire protein with large *C*^*j*^ values (Fig. 4). These include 5 residues on the α-helical N-terminus and 27 residues on the β-barrel. Experimental evidence also indicates that both the N-terminal α-helix and the β-barrel contribute to voltage sensitivity (64, 65, 81, 82). Together with our data, this suggests a scenario where the entire protein is involved in the response to changes in the MP rather than a single domain playing the role of voltage sensor. Among the residues pinpointed by our analysis, mutagenesis of D16, K20, K61 and E84 is known to modulate the steepness of the current-voltage relationship (64), while that of D32 and K96 does not significantly affect voltage sensitivity (64). This indicates there are likely a few false positive signals in our analysis; therefore it has to be combined with experiments to confirm voltage sensitivity of the detected elements.

Previous experimental studies performed on ClC1 have revealed two sources of voltage sensitivity in this protein. Mutagenesis of the so-called gate residue E148 completely abolishes voltage sensitivity (29, 97, 102), and changes in Cl-concentration affects gating charge responsible for activation (66, 68–70). Our analysis shows large *C*^*j*^ for both E148 and Cl-ions bound along the conduction path (Fig. 5). Moreover, it also pinpoints a region at the interface between the two ClC1 subunits; in particular, K147, K165, K409, E225 and E490 all show *C*^*j*^ above the detection threshold, suggesting they are likely significant contributors to the ClC1 gating charge.

In the case of the M_2_ receptor, the identified voltage-sensing element is complex and composed of the protein residues and a Na^+^ ion bound inside the internal hydrophilic pocket. The role of a Na^+^ ion in M_2_ receptor voltage sensitivity was in fact recently suggested based on molecular dynamics simulations (67), while mutagenesis experiments revealed a crucial role of D69 in generation of the gating current (84). Our data agrees with these findings and shows large *C*^*j*^ for D69, a Na^+^ ion and also for D107 (Fig. 5). In addition, our results support the recent experimental finding that the DRY motif on the TM3 segment does not contribute to voltage sensitivity (84).

Finally, for the Na^+^/K^+^ ATP-ase, we found three regions where *C*^*j*^ is above the detection threshold, including the ionic binding site, the extracellular and cytosolic openings (Fig. 5). The ionic binding site, which is composed of E327, D808, D926, E953 and three Na^+^ ions, shows the largest contribution to the gating charge. This agrees with the known data reporting that changes in the extracellular Na^+^ concentration has a strong effect on voltage sensitivity of the Na^+^/K^+^ ATP-ase current (8, 9, 71, 72).

## Conclusions

One of the reasons voltage sensitivity in membrane proteins is hard to detect is that it depends on a combination of two factors: the ability of a protein to focus an electric field in some of its regions and its ability to respond to the electric field based on the presence of charged elements in these regions. We show that it is possible to use a simple and cheap computational tool to distinguishing between voltage-sensitive and voltage-insensitive proteins, and to detect true voltage-sensitive residues in voltage-gated ion channels. The application of these tools to six membrane proteins without well-characterized voltage sensors leads to the prediction of several candidates for voltage-sensing elements, some of which have already been shown to contribute to voltage sensitivity while others remain to be tested experimentally. The suggested approach is general, it does not require extensive computational studies, and it can be applied to any other membrane protein regardless of structure or function. This application is important to unravel the variety of ways voltage sensitivity has been manifested in biological molecules, and it should be highly useful both to identify how different factors such as mutations, membrane composition, or ligand binding might alter voltage sensitivity and to engineer this property into voltage-insensitive membrane proteins.

## Methods

### Connection between *C*^*j*^ and *Q*^*j*^

Equation (2), which we use to estimate the capacity of the system’s element *j* to respond to changes in the local electric field *C*^*j*^, can be rewritten as:

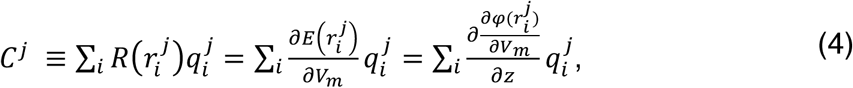

where 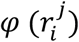 is the local electrostatic potential measured at the position 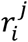, and the summation runs over all charges 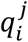 of the element *j*. The numerator in Eq. (4) 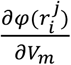 can be substituted for the coupling function 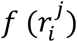, which by definition represents the coupling of the charge 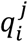 to the local electrostatic potential and has a direct connection to the gating charge *Q*^*j*^ (103):

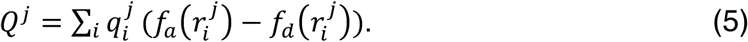

Here, *Q*^*j*^ is represented as a charge-weighted difference between the coupling functions of the activated and deactivated states 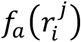 and 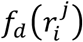, respectively. Combining Eqs. (4) and (5) we obtain:

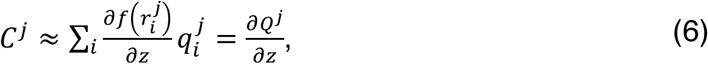

where 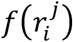 corresponds to a conformational state for which the high-resolution structure has been obtained. Therefore, *C*^*j*^ is equivalent to *Q*^*j*^ when the residue *j* is displaced by 1 Å along the direction of the external electric field *z*.

### Systems’ preparation and Molecular Dynamics simulations

The Charmm-GUI server was used to prepare the systems for molecular dynamics simulations (104). Briefly, every protein of interest was embedded into a 1-palmytoyl-2-oleoylphosphatidylcholine (POPC) bilayer and solvated with 150 mM of either KCl or NaCl solution. The CHARMM36 force field (105) was used to describe proteins and lipids with TIP3P as a water model (106). Note that the Na^+^ ion bound inside the protein cavities of the M_2_ receptor was not placed there initially; instead, it reached the binding site during the molecular dynamics simulations. For the detailed description of the systems composition and properties, see Table 2.

**Table 2.**

| Protein | PDB code | Residue patching | System's dimensions [ÅxÅxÅ] | Number of atoms | Duration of the equilibration last step |
| --- | --- | --- | --- | --- | --- |

|  |  |  |  |  | [ns] |
| --- | --- | --- | --- | --- | --- |
| Shaker* | A homology model based on 2r9r (90) see (89) for the details | none | 130x130x105 | 191.034 | 1 μs |
| NavMs | 5hvx (91) | none | 145x145x120 | 209.838 | 400 ns |
| GLIC | 4npq (100) | none | 110x110x145 | 159.083 | 1 μs |
| TWIK-1 | 3ukm (101) | disulfide bond between C78 and C78 (intersubunit) | 105x105x125 | 127.219 | 200 ns |
| Cx26 | 2zw3 (92) | disulfide bonds between C45 and C156, C50 and C152, C56 and C145 | 130x130x122 | 189.168 | 200 ns |
| VDAC1 | 3emn (96) | none | 85x85x85 | 54.590 | 200 ns |
| TRPV1** | 3j5r (93–95) | none | 160x140x175 | 402.346 | 750 ns |
| CIC1 | 1ots (97) | none | 130x130x100 | 158.553 | 200 ns |
| M<sub>2</sub> receptor | 3uon (98) | disulfide bonds between C96 and C176, C413 and C416 | 90x90x105 | 76.169 | 200 ns |
| Na<sup>+</sup>/K<sup>+</sup> ATP-ase*** | 3wgu (99) | D195, E358, E779, D804, E954 are protonated; disulfide bonds between C126 and C149, C159 and C175, C213 and C276 | 115x115x190 | 250.192 | 100 ns |

\* The trajectory used for the analysis was taken from (89).
\*\* The trajectory for the closed capsaicin-bound state of TRPV1 used for the analysis was taken from (95).
\*\*\* The trajectory for the sodium-bound Na<sup>+</sup>/K<sup>+</sup> ATP-ase used for the analysis was taken from (107).

The molecular dynamics simulations were performed using GROMACS 2016.1 (108). Each system was equilibrated following a multi-step protocol. During the first 2.5 ns, the protein and lipid headgroups were restrained to their initial positions to allow for the rearrangements of lipid tails and solution. The lipid headgroups were subsequently released and the simulations continued for 2.5 ns. In the next 35 ns, the restraints applied to the protein backbone and sidechains were gradually decreased from 4000 kJ/mol/nm and 2000 kJ/mol/nm, respectively, to 0 kJ/mol. Finally, the protein was fully relaxed without restraints until the root mean square deviation with respect to the initial structure reached a plateau value (for the duration of this last step of the equilibration, see Table 1). The final frame of the equilibration trajectory was used for the simulations under an electric field.

A Nose-Hoover thermostat (109) and Parrinello-Rahman barostat (110) were used to keep temperature (300 K) and pressure (1 atm) constant. A cutoff of 1.2 nm was applied for short-range electrostatics and VdW interactions. For the latter, a switching function between 1.0 and 1.2 nm was applied to smoothly bring the forces to 0 at 1.2 nm. Particle Mesh Ewald summation (111) was used for the long-range component of electrostatics. 1 fs time step was used for the first two steps of the equilibration, and 2 fs – for the rest of the equilibration and the simulations under an electric field.

### Molecular Dynamics simulations under an electric field

For each protein of interest, 10 conformational states were extracted from the equilibration trajectory with a stride of 20 ns. For every state, 9 short (2 ns) molecular dynamics simulations were performed with the restraints applied to all heavy atoms of the protein and under an electric field; the following values of the field were used: −0.08, −0.06, −0.04, −0.02, 0, 0.02, 0.04, 0.06 and 0.08 V/Å. 5 frames were extracted from each of the trajectories to compute an average local electrostatic potential map *φ* using the PMEpot plugin of VMD (112):

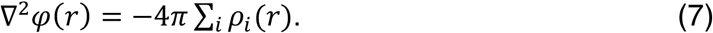

Here, *ρ*_*i*_(*r*) corresponds to a point charge approximated by a spherical Gaussian with an Ewald factor of 0.25. Equation 7 was solved on a grid with a resolution of 1 Å. *R*, *R*^*j*^ (per-residue local electric field response: 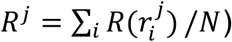, and *C*^*j*^ were calculated using Eqs. (1) and (2) with an in-house python code. Finally, the average and the standard deviations of *R*^*j*^ and *C*^*j*^ were computed based on all 10 conformational states and also all subunits in the case of homo-multimers.

The detection threshold was identified based on the calculations of *R*^*j*^ (Fig. S1). For every protein of interest *R*^*j*^ was computed for the two dimensions orthogonal to that of the external electric field (*i.e.* in the Eq. (1) the components of the local field orthogonal to the external field were used). The largest *R*^*j*^ value, 0.02 Å^-1^, was further considered as the detection threshold.

## Acknowledgments

The simulations were performed on resources provided by the Swedish National Infrastructure for Computing (SNIC) at PDC Centre for High Performance Computing (PDC-HPC). This work was supported by grants from the Swedish Research Council (2017-04641) and Science for Life Laboratory. We acknowledge Kawai Lee, Marie Lycksell and Rebecca J. Howard for providing the GLIC trajectory, Asghar Razavi for providing the Na^+^/K^+^ ATP-ase trajectory, and Samira Yazdi for providing the Shaker trajectory. The authors have declared that no competing interests exist.

## Author contributions

M.A.K., E.L. and L.D. have designed the research. M.A.K. has performed the simulations, analyzed the data and written the original draft. M.A.K., E.L. and L.D. have edited the paper.

